# CoV-Seq: SARS-CoV-2 Genome Analysis and Visualization

**DOI:** 10.1101/2020.05.01.071050

**Authors:** Boxiang Liu, Kaibo Liu, He Zhang, Liang Zhang, Yuchen Bian, Liang Huang

## Abstract

**Summary:** COVID-19 has become a global pandemic not long after its inception in late 2019. SARS-CoV-2 genomes are being sequenced and shared on public repositories at a fast pace. To keep up with these updates, scientists need to frequently refresh and reclean datasets, which is ad hoc and labor-intensive. Further, scientists with limited bioinformatics or programming knowledge may find it difficult to analyze SARS-CoV-2 genomes. In order to address these challenges, we developed CoV-Seq, a webserver to enable simple and rapid analysis of SARS-CoV-2 genomes. Given a new sequence, CoV-Seq automatically predicts gene boundaries and identifies genetic variants, which are presented in an interactive genome visualizer and are downloadable for further analysis. A command-line interface is also available for high-throughput processing.

**Availability and Implementation:** CoV-Seq is implemented in Python and Javascript. The webserver is available at http://covseq.baidu.com/ and the source code is available from https://github.com/boxiangliu/covseq.

**Supplementary information:** Supplementary information are available at *bioRxiv* online.

## 1 Introduction

The 2019 Novel Coronavirus (SARS-CoV-2) caused an outbreak of viral pneumonia since late 2019 and has become a global pandemic. Despite efforts to contain its spread, SARS-CoV-2 has infected more than 4 millions patients and caused nearly 300,000 deaths worldwide as of early May (Dong *et al.*, 2020). To understand its evolution and genetics, scientists have sequenced SARS-CoV-2 genomes from patients across different age groups, genders, ethnicities, locations, and disease stages (Hadfield *et al.*, 2018). These genomic sequences are being shared on public repositories at a rapid pace, with thousands of new sequences every week (Elbe and Buckland-Merrett, 2017; Brister *et al.*, 2015; Kanz *et al.*, 2005; Wei, 2019). To keep up with the latest developments, scientists need to frequently download and clean new datasets, which is ad hoc and time-consuming. On the other hand, scientists with limited knowledge in bioinformatics or programming may experience difficulty in analyzing SARS-CoV-2 genomes.

We developed the CoV-Seq toolkit to address these challenges. CoV-Seq consists of several components: a data analysis pipeline that takes FASTA sequences and generates variant callsets in VCF format and open reading frame (ORF) predictions. The pipeline automatically filters low-quality sequences and removes duplicate sequences, performs sequence alignment, as well as identifies and annotates genetic variants (Figure 1A). We provide a webserver (http://covseq.baidu.com/) to allow rapid analysis of custom sequences without any programming. The web interface includes an interactive genome visualizer and tabulated displays of genetic variants and ORF predictions. All results can be downloaded for downstream analysis. Further, we provide a command-line interface to allow high-throughput processing on local environments. To facilitate data sharing, we aggregate SARS-CoV-2 sequences from GISAID (Elbe and Buckland-Merrett, 2017), NCBI (Brister *et al.*, 2015), EMBL (Kanz *et al.*, 2005) and CNGB (Wei, 2019), and publish annotated variant callsets and metadata on a weekly basis.

**Fig. 1.**
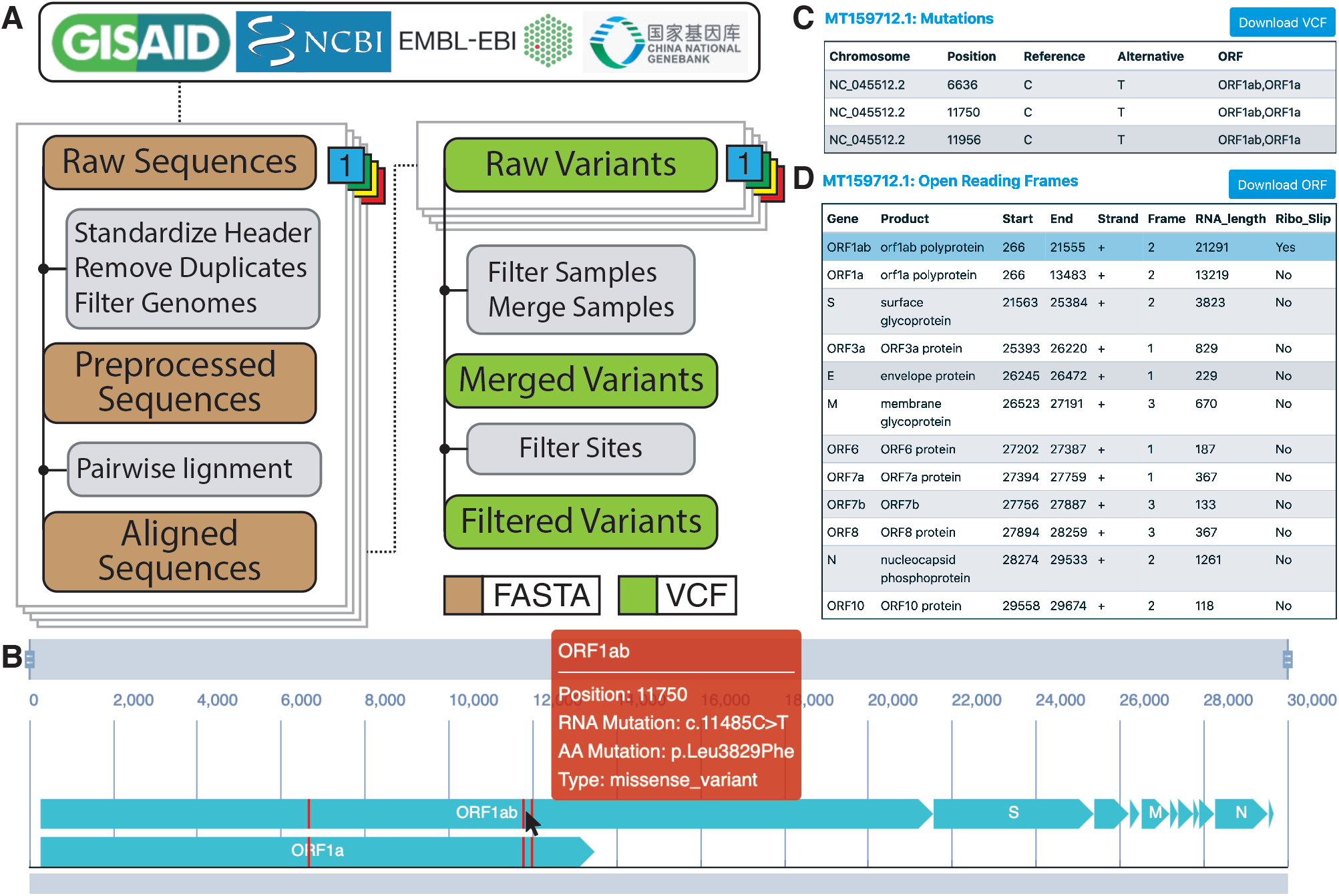
The CoV-Seq pipeline and web interface. (A) Genomic sequences are collected from GISAID, NCBI, EMBL and CNGB. We remove incomplete genomes (length < 25,000) and duplicate genomes before alignment with MAFFT against the reference genome NC_045512.2. We use a custom Python script to generate raw variant calls and remove samples with too many mutations indicative of sequencing error. After merging VCFs, we remove multi-allelic sites and variants with the poly-A tail for a filtered set of variants. (B) The interactive genome visualizer shows ORFs (turquoise) and mutations (red). Users can zoom with the top bar and pan with the bottom bar. Hovering over gene bodies and mutations will trigger pop-up windows for relevant information. (C) The mutation table shows positions, alleles and intersecting ORFs. (D) The ORF table shows predicted gene boundaries and supporting information.

## 2 Analysis and Visualization of Viral Genomes

The collection of SARS-CoV-2 genomic sequences is rapidly expanding. An integrated pipeline is essential for keeping current with frequent updates. To understand the evolution and genetics of a new viral strain, it is necessary to identify its genetic mutations and gene boundaries. Existing software packages such as VAPiD (Shean *et al.*, 2019) and VIGOR (Wang *et al.*, 2010) focus on gene annotation. To our knowledge, a software package to identify, annotate, and visualize genetic variants for SARS-CoV-2 is lacking.

CoV-Seq provides an intuitive web interface for analyzing and visualizing SARS-CoV-2 variants and ORFs. Upon receiving custom sequences, CoV-Seq identifies gene boundaries and genetic variants and displays them alongside a full-length genome (Figure 1B). Users interact with the genome by dragging the zoom bar to adjust the magnification and the position bar to pan along the genome. When the genome window contains less than 150 nucleotides, letters will appear to indicate both the nucleotide bases and amino acid residues. Hovering over gene bodies or variants will trigger pop-up windows for relevant information (Figure 1B red box). Clicking on an ORF brings up three tables. A first table shows all variants and their positions, alleles, and ORFs in which they belong (Figure 1C). A second table shows ORF annotations obtained by aligning input sequences against the reference sequence and transferring annotations from Genbank (Figure 1D). A third table shows nucleotide and protein sequences for the selected ORF. All tables can be downloaded for further analysis.

## 3 SARS-CoV-2 genome processing pipeline

Extraction of genetic variants from sequences involves the following steps (Figure 1A): (1) preprocessing: removal of low-quality and duplicated sequences; (2) alignment: pairwise alignment of each sequence against a reference; (3) variant calling: identify differences between aligned sequences; and (4) postprocessing: remove low-quality sites and annotate variants.

We aggregate SARS-CoV-2 sequences from GISAID, NCBI, EMBL and CNGB. Many sequences represent incomplete genomes, sometimes containing only a single gene. We filter these genomes using a lenient cutoff of 25,000 nucleotide because it removes distinctly incomplete genomes while retains complete genomes (Figure S1). Both NCBI and EMBL are part of the International Nucleotide Sequence Database Collaboration (INSDC) and therefore contain duplicate submissions, which we remove by comparing the Accession IDs. Further, dual submissions can appear in both GISAID and INSDC under different Accession IDs. We consider two submissions as suspect duplications if they have identical genomic sequences. These suspect duplications are marked in the metadata but not removed because it is possible for an identical strain to infect multiple patients. We perform pairwise alignment against the reference sequence NC_045512.2 using MAFFT (Katoh and Standley, 2013) and use a custom Python script for variant calling. We left-normalize each variant with bcftools (Li, 2011) and remove samples with too many variants indicative of sequencing error. We use a lenient cutoff of 150 variants because it removes samples with extremely large numbers of variants while keeping most samples (Figure S2). During postprocessing, we remove multi-allelic sites because these sites are more likely to occur in regions prone to sequencing error, such as the two ends of the genome (Figure S3). Further, we remove variants within the poly-A tail. Filtered variant callset is annotated with snpEff (Cingolani *et al.*, 2012). Both raw and filtered VCF files are available for download on CoV-Seq (http://covseq.baidu.com/browse) and the pipeline is open source at https://github.com/boxiangliu/covseq. We are committed to update VCF files and associated metadata weekly on Mondays.

## Acknowledgements

We thank the Chinese Center for Disease Control and Prevention for co-developing the idea and Xing Li for helpful discussions.

## Notes

http://covseq.baidu.com/

### Competing Interest Statement

The authors have declared no competing interest.

