## Supplemental Material for "CoV-Seq: SARS-CoV-2 Genome Analysis and Visualization"

May 11, 2020

### 1 Supplementary Figures

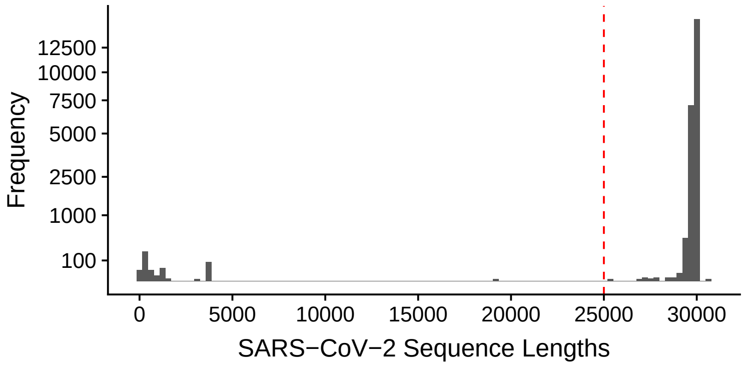


Figure S1: Distribution of SARS-CoV-2 sequence lengths. Sequences with lengths less than 25,000 nucleotides are removed during preprocessing.

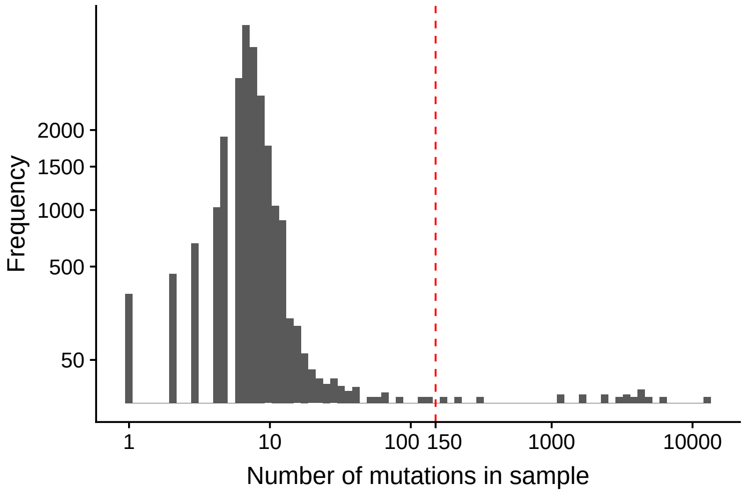


Figure S2: Distribution of sample mutations identified against the reference genome NC_045512.2. Samples with more than 150 mutations are removed during post-processing.

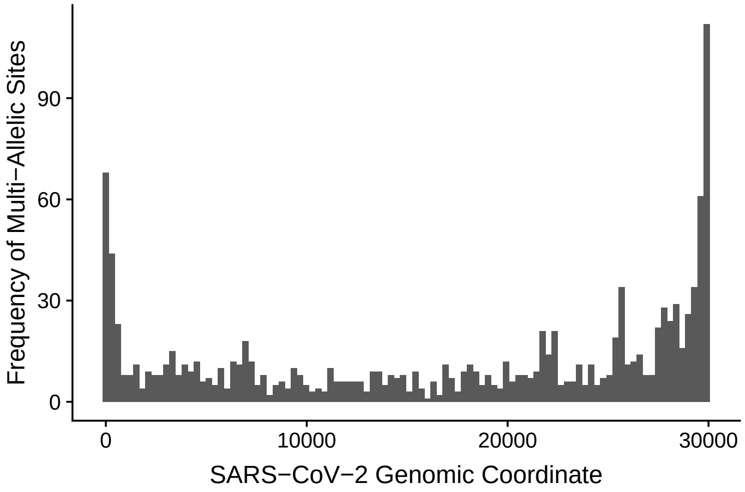


Figure S3: Distribution of multi-allelic sites along the SARS-CoV-2 genome. Multi-allelic sites are more likely to be identified at the beginning and the end of the genome.
